# A variant selection framework for genome graphs

**DOI:** 10.1101/2021.02.02.429378

**Authors:** Chirag Jain, Neda Tavakoli, Srinivas Aluru

## Abstract

**Motivation:** Variation graph representations are projected to either replace or supplement conventional single genome references due to their ability to capture population genetic diversity and reduce reference bias. Vast catalogues of genetic variants for many species now exist, and it is natural to ask which among these are crucial to circumvent reference bias during read mapping.

**Results:** In this work, we propose a novel mathematical framework for variant selection, by casting it in terms of minimizing variation graph size subject to preserving paths of length *α* with at most *δ* differences. This framework leads to a rich set of problems based on the types of variants (SNPs, indels), and whether the goal is to minimize the number of positions at which variants are listed or to minimize the total number of variants listed. We classify the computational complexity of these problems and provide efficient algorithms along with their software implementation when feasible. We empirically evaluate the magnitude of graph reduction achieved in human chromosome variation graphs using multiple *α* and *δ* parameter values corresponding to short and long-read resequencing characteristics. When our algorithm is run with parameter settings amenable to long-read mapping (*α* = 10 kbp, *δ* = 1000), 99.99% SNPs and 73% indel structural variants can be safely excluded from human chromosome 1 variation graph. The graph size reduction can benefit downstream pan-genome analysis.

**Implementation:** https://github.com/at-cg/VF

## 1 Introduction

High-throughput technologies enable rapid sequencing of numerous individuals in a species population and cataloging observed variants. This is leading to a switch from linear representation of a chosen reference genome to graph representations depicting multiple observed haplotypes. Graph representations more accurately reflect the sampled individuals within a population, and their use in genome mapping algorithms reduces reference bias and increases mapping accuracy when sequencing a new individual (Ballouz *et al*., 2019). There is abundant research on data structures designed for graph representations of genomes and pangenomes (Garrison *et al*., 2018; Li *et al*., 2020), their space-efficient indexing (Sirén *et al*., 2014; Marcus *et al*., 2014; Holley *et al*., 2016; Ghaffaari and Marschall, 2019; Jain *et al*., 2019b; Chang *et al*., 2020; Kuhnle *et al*., 2020), and alignment algorithms (Kuosmanen *et al*., 2018; Jain *et al*., 2020; Rautiainen and Marschall, 2020; Darby *et al*., 2020; Ivanov *et al*., 2020) to map sequences to reference graphs. For review papers summarizing these developments, see (Paten *et al*., 2017; Computational Pan-Genomics Consortium, 2018; Eizenga *et al*., 2020).

While graph representations have numerous advantages, complete variation graphs that include every variant have certain drawbacks. The graphs invariably contain paths combining variants across haplotypes, but never seen in any observed haplotype. The number of such recombinant paths increases combinatorially with graph size, and is particularly troublesome when mapping long reads which span greater distances. Accuracy of sequence-to-graph mapping algorithms shows diminishing returns at larger graph sizes, and is even negatively affected eventually (Pritt *et al*., 2018; Sirén *et al*., 2020). The computational complexity of the mapping algorithms makes it impractical to apply them at the scale of complete variation graphs. A few attempts have been made to address the first issue by augmenting paths with haplotype information and specifically developing haplotype-aware indexing strategies (Iqbal *et al*., 2012; Sirén *et al*., 2020; Mokveld *et al*., 2020).

The aforementioned factors point to the need for variant selection algorithms which tame reference graph sizes, and strike the right balance for subsequent mapping accuracy and speed. This was primarily approached through selecting variants from a specific database (Schneeberger *et al*., 2009; Danek *et al*., 2014; Liu *et al*., 2016), based on allelic frequency (Maciuca *et al*., 2016; Eggertsson *et al*., 2017; Kim *et al*., 2018), or specific to a biological context such as limiting to a particular population (Sirén *et al*., 2014) or genome loci of interest (Vijaya Satya *et al*., 2012; Dilthey *et al*., 2015; Jain *et al*., 2019a). Recently, Pritt *et al*. (2018) developed a more systematic approach FORGe by developing a mathematical model to prioritize variants, and selecting top scoring variants according to the model. In FORGe, the ranking of each variant is done based on its frequency in a population, and its contribution to runtime and space overhead of a read-to-graph mapper.

In this work we propose a rigorous algorithmic framework for variant selection from the perspective of preserving subsequent mapping accuracy. Consider a complete variation graph constructed from a set of given haplotypes. Any substring of a haplotype has a corresponding path in the complete variation graph. Not including some variants will introduce errors in the corresponding paths. If the number of such errors is matched with the error tolerance built into sequence-to-graph mapping algorithms, the same identical paths can still be found. We make the following contributions:

- We develop a novel mathematical framework for variant selection subject to preserving paths of length *α* while allowing at most *δ* differences. We separately consider the problems of minimizing the number of positions at which variants are retained, and minimizing the total number of variants selected.
- We show that both problems are optimally solvable in polynomial time when only SNPs are considered and the goal is to preserve all paths of length *α* found in the complete variation graph.
- These problems become challenging when indels are brought into play. We present efficient heuristics that guarantee preserving paths of length *α* while allowing at most *δ* edits, but do not guarantee optimal reduction in graph size.
- We empirically evaluate run-time performance and reduction in variation graph sizes achieved by the multiple algorithms that are proposed in this paper. For testing, we utilize human chromosome sequences, SNP variants from the 1000 Genomes Project (Consortium *et al*., 2015), and indel structural variants (SVs) (i.e., indels of size ≥50 bp) from fifteen diverse humans (Audano *et al*., 2019). When chromosome 1 variation graph is built using SNP variants, and parameters amenable to short reads (*α* = 150 and *δ* = 8) are used, the reduced graph excludes 94.44% SNPs. With parameters adjusted for long reads (*α* = 10 kbp and *δ* = 1000), 99.99% SNPs are excluded. When SVs are considered, the (*α* = 150 bp, *δ* = 8) and (*α* = 10 kbp, *δ* = 1000) settings result in excluding 0% and 73% SVs respectively.
- Finally, we consider the complexity of haplotype-aware versions of the above problems where the goal is to only preserve paths of length *α* actually found in the input haplotypes (i.e., not recombinant paths), and prove that they are 𝒩𝒫-hard even for *δ* = 1.

## 2 Proposed framework

Let *R*_1_, *R*_2_, …, *R*_*m*_ be *m* input reference haplotype sequences. To be consistent with current literature, we assume one of these (say *R*_1_) is a special reference and the other haplotypes are described as variations from it. A (complete) variation graph of these sequences is represented using an edge-labeled directed multigraph *G*(*V, E, σ*) as follows. The graph consists of haplotype *R*_1_ as a linear backbone, augmented with the set of variants present in *R*_2_, *R*_3_, …, *R*_*m*_, assumed to be known *a priori*. Each variant represents a deviating base from *R*_1_ (SNP) or an insertion/deletion (can be multiple bases). The function *σ* : *E* → Σ ∪ {*ϵ*} specifies edge labels, where Σ denotes the alphabet and *ϵ* denotes the empty character. The haplotype *R*_1_ is represented in *G* as a directed chain with character labeled edges such that the chain spells the sequence *R*_1_. This chain will have |*R*_1_| + 1 ordered vertices 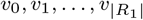. These vertices serve as a convenient *coordinate axis* for the variation graph. Each SNP variant is an additional labeled edge between vertices at two adjacent coordinates. A deletion variant is an edge labeled *ϵ* between a pair of vertices, whose coordinates are separated by the deletion length. An insertion variant is represented as a chain of one or more labeled edges that starts and ends at the same vertex. In this setup, the total number of variants at coordinate *i* (0 ≤ *i* ≤ |*R*_1_|) equals out-degree of the vertex *v*_*i*_ minus one. See Figure 1 for an illustration.

**Fig 1.**
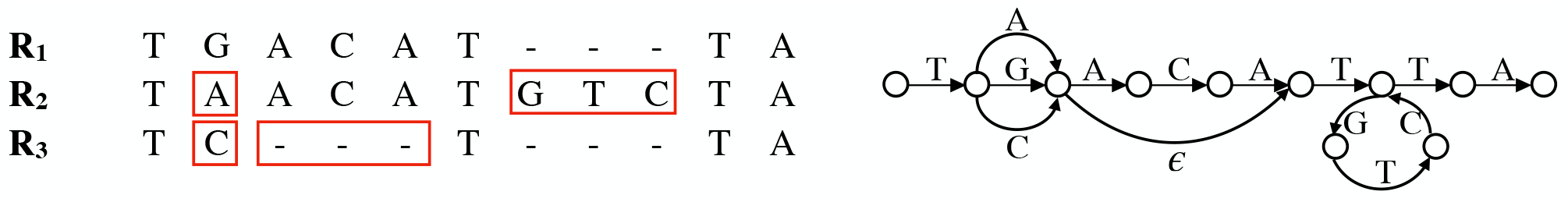
An example to illustrate construction of variation graph from three haplotype sequences.

Any path in graph *G* with *α* non-empty edges spells a string of length *α*. We place the restriction that a path is allowed to visit a vertex at most twice. This restriction avoids traversal of more than one insertion variant at the same coordinate. Note that any recombination of variants that occur at different positions is allowed. Thus, the graph contains paths corresponding to each haplotype and any substrings thereof, but also numerous additional paths (genotypes) that are not present in any haplotype. It is unknown whether such a recombinant genotype exists in the population or not. Restricting paths to only those that belong to at least one input haplotype can also be useful, and will be considered separately (Section 4).

We seek to compute a reduced variation graph *G′* (*V ′, E ′, σ ′*), where *V ′* ⊆ *V, E ′* ⊆ *ϵ*, and for all *e* ∈ *E ′, σ ′* (*e*) = *σ*(*e*). The reduced graph *G′* corresponds to removing some variants in graph *G*. Our goal is to reduce graph *G*(*V, E, σ*) to the maximum extent possible while ensuring that any *α*-long string corresponding to a path in *G* can be mapped to the same starting vertex (coordinate) in *G′* without exceeding a user-specified error threshold *δ*. In practice, *α* should be a function of read lengths whereas *δ* is determined based on sequencing errors and error-tolerance of read-to-graph mapping algorithms.

We formulate four versions of the problem based on what types of variants are allowed and the reduction objective. First consider the case where all variants are SNPs.

### Definition 1.

*Graph G′ is said to be* (*α, δ*)_*h*_*-compatible if all α-long strings in graph G can be mapped to their corresponding paths in graph G′ with Hamming distance* ≤ *δ*.

### Problem 1.

*Compute an* (*α, δ*)_*h*_*-compatible reduced variation graph G′ with minimum number of coordinates containing one or more variants*.

### Problem 2.

*Compute an* (*α, δ*)_*h*_*-compatible reduced variation graph G′ with minimum number of variants*.

In Problem 1, we seek to ‘linearize’ the graph, whereas in Problem 2, we intend to remove as many variants as possible. A user can choose either version based on downstream analysis. For the next two problem versions, suppose the variant set also contains indels.

### Definition 2.

*Graph G′ is said to be* (*α, δ*)_*e*_*-compatible if all α-long strings in graph ′can be mapped to their corresponding paths in graph G′ with edit distance* ≤ *δ*.

### Problem 3.

*Compute an* (*α, δ*)_*e*_*-compatible reduced variation graph G′with minimum number of coordinates containing one or more variants*.

### Problem 4.

*Compute an* (*α, δ*)_*e*_*-compatible reduced variation graph G′with minimum number of variants*.

In Problems 1 and 2 that consider only SNP variants, *α*-long paths will begin at a vertex along the coordinate axis as there are no other vertices introduced in the graph. In Problems 3 and 4 however, a path can also begin at other vertices due to insertion variants. In this case, we assume an *α*-long string that maps to *G* must also be mappable starting from the corresponding vertex in *G′* if that insertion variant is preserved. If the variant is not preserved, it must be mappable to the closest vertex along the coordinate axis.

## 3 Proposed algorithms

### 3.1 Results for variation graphs with SNPs

#### 3.1.1. Greedy algorithm for Problem 1

As the goal is to minimize the number of coordinates (positions along the special reference *R*_1_) at which variants are included in *G′*, we either fully *remove* or fully *retain* all the variants at each coordinate with variants. When removing variants at a coordinate, its outgoing edge label is chosen to be the base from *R*_1_. However, the (*α, δ*)_*h*_-compatibility is sustained even if the base is chosen from a different haplotype, or any arbitrary character in Σ.

A path of length *α* naturally corresponds to a line segment of length *α* starting at an integer coordinate. Observe that in any *α*-long segment, we cannot remove variants at *> δ* coordinates without violating the (*α, δ*)_*h*_-compatibility of reduced graph *G′* (Figure 2(a)). A variant coordinate *i* is contained in *α* segments of length *α* each, whose starting positions are in [*i* − *α* + 1, *i*]. For each variant position, we associate two events with coordinates *start*_*i*_ = *min*{0, *i* − *α* + 1} and *end*_*i*_ = *i* respectively. Assuming that the *n* SNP coordinates are given as sorted array, the corresponding 2*n* events can be sorted in *O*(*n*) time. When two events have equal coordinates, the *start* event type should be placed earlier than the *end* event type in the sorted order.

**Fig 2.**
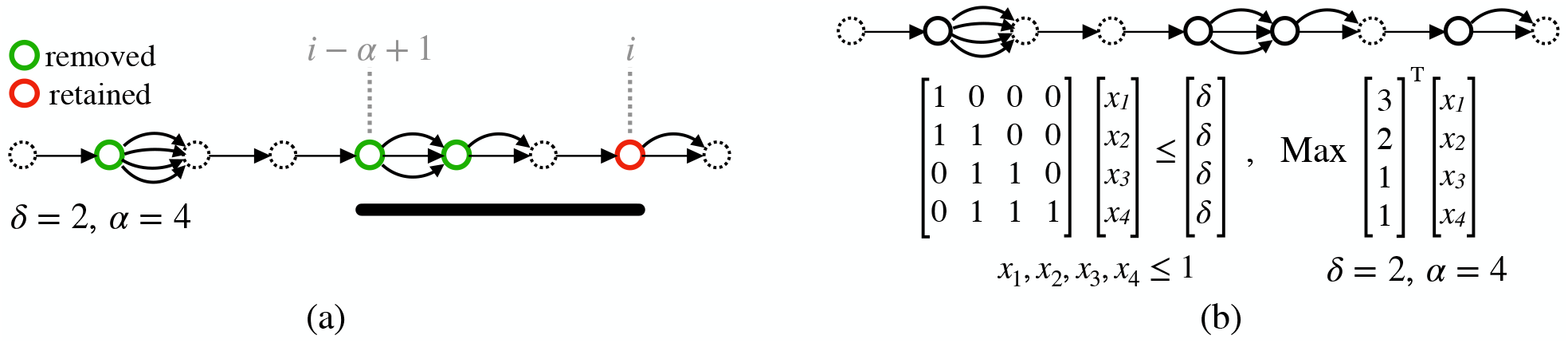
Execution of the Greedy and LP algorithms on an example variation graph containing SNPs only. Edge labels are not shown as they do not affect the execution of either algorithm.

Our greedy algorithm works as follows. Begin by placing an *α*-long segment at position 0, and remove variants in the leftmost *δ* variant positions and retain the rest (if any). Keep a count of the number of positions within the current segment at which variants are removed. Iteratively consider each event in the sorted order. If the event is of type *start*_*i*_ and the count is less than *δ*, the variants at position *i* are removed and the count is incremented by one. If the event is of type *start*_*i*_ but the count is equal to *δ*, the variants at position *i* are retained. If the event is of type *end*_*i*_ and the variants at *i* were previously removed, the count is decremented by 1. As can be seen, the algorithm maintains (*α, δ*)_*h*_-compatibility and runs in *O*(*n*) time.

##### Proof of optimality

Suppose the greedy algorithm retains variants at coordinates *g*_1_, *g*_2_, …, *g*_*p*_ in ascending order. Let *o*_1_, *o*_2_, …, *o*_*q*_ be the ordered variant coordinates retained by an optimal solution. Let *k* be the first position where the solutions differ, i.e., *g*_*j*_ = *o*_*j*_ for *j < k* and *g*_*k*_ ≠ *o*_*k*_. Due to our greedy strategy, *o*_*k*_ *< g*_*k*_. Though *o*_*k*_ was chosen by the optimal algorithm, (*α, δ*)_*h*_-compatibility is not violated until start event for *g*_*k*_ is reached. For any path starting at a later coordinate, retaining variants at *g*_*k*_ offers the same benefit as retaining at *o*_*k*_. Thus, replacing *o*_*k*_ with *g*_*k*_ will maintain optimality and (*α, δ*)_*h*_-compatibility. Hence, the greedy solution is also optimal.

###### Lemma 1.

*The above greedy algorithm solves Problem 1 in O*(*n*) *time*.

#### 3.1.2 A linear programming solution to Problem 2

Here, we seek to minimize the total number of variants retained. Interestingly, we can show that optimal solutions still retain or remove all variants at a coordinate.

##### Lemma 2.

*An optimal solution to Problem 2 either retains or removes all variants at a coordinate*.

##### Proof.

By contradiction. Suppose there exists an optimal reduced graph *G′* with partially removed variants at coordinate *i*. Coordinate *i* already induces an error in some *α*-long paths in *G* that contain the coordinate. Accordingly, removal of all variants at coordinate *i* can be tolerated by all *α*-long paths containing that coordinate, further implying that graph *G* must be sub-optimal.

Suppose *G′* contains no variants at coordinate *i*, then this choice reduces the count of variants by *out*(*v*_*i*_) − 1, where *out*(*v*_*i*_) is the out-degree of vertex *v*_*i*_. As can be seen, Problem 2 is harder than Problem 1 because the number of variants at different coordinates can be different, leading to variable gains. We address this problem by using an Integer Linear Programming (ILP) system that is polynomially-solvable using LP relaxation.

Let *p*_1_, *p*_2_, …, *p*_*n*_ be the *n* variant coordinates in *G* in ascending order. Let *X* be an *n* × 1 boolean column vector where *X*[*i*] = 1 iff variants are removed at coordinate *p*_*i*_ in creating *G*. ′ Let *C* be another *n*×1 column vector where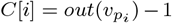, i.e., the reduction achieved in variant count by removing variants at *p*_*i*_. The goal is to maximize *C*^*T*^ *X*. Next, we specify constraints to ensure (*α, δ*)_*h*_-compatibility of graph *G*, ′ by not allowing removal of variants at *> δ* coordinates in any *α*-long segment. Similar to the observation made while addressing Problem 1, it suffices to check this constraint only in the subset of *α*-sized segments that end at the *n* variants. Accordingly, let *A* be a boolean *n* × *n* matrix such that *A*[*i*][*j*] = 1 iff coordinate *p*_*j*_ is within the *α*-sized segment range (*p*_*i*_ − *α, p*_*i*_] of coordinate *p*_*i*_. Then, ILP constraints required to ensure (*α, δ*)_*h*_-compatibility of *G′* can be specified as *A* · *X* ≤ *B*, where *B* is an *n* × 1 column vector with each value = *δ*. We also need to ensure that the *X*[*i*]’s are boolean. This can be achieved by expanding *A* to a 2*n* × *n* matrix with the bottom *n* rows being the *n* × *n* identity matrix, and similarly expanding *B* to a 2*n* × 1 vector with the bottom *n* entries set to 1. Now, maximizing *C*^*T*^ *X* while satisfying *A* · *X* ≤ *B* leads to an optimal reduced graph *G′* that is (*α, δ*)_*h*_-compatible.

##### Run-time complexity

Matrix *A* exhibits a special structure that guarantees integral optimal LP solutions. Observe that *A* is a 0-1 matrix, and the 1’s appear consecutively in each row of *A* which makes it an interval matrix (Fulkerson and Gross, 1965). As a result, the above ILP can be solved in polynomial time by solving the corresponding LP, which has *O*(*n*^*ω*^) run-time complexity where *ω* is the exponent of matrix multiplication (van den Brand, 2020).

###### Lemma 3.

*The above LP-based algorithm solves Problem 2 in O*(*n*^*ω*^) *time*.

### 3.2 Results for variation graphs with SNPs and indels

Variation graphs with indels introduce additional complexity. When considering only SNPs, we benefited from the fact that end vertices of any *α*-long paths will be located on the coordinate axis. In addition, right end of a path was a fixed distance away from its left along the coordinate axis. When indels are permitted, these properties are no longer true, making Problems 3 and 4 more challenging. We present two heuristic solutions, each of which can be used to solve either problem.

#### 3.2.1. A greedy algorithm

We first propose a ‘conservative’ greedy heuristic which guarantees an (*α, δ*)_*e*_-compatible reduced graph that is not necessarily optimal in terms of the desired reduction objectives. We choose to either retain or remove all variants at a coordinate vertex (vertex along *R*_1_). Suppose a coordinate vertex *v* has all three types of variants, i.e., insertions, deletions and SNPs. We evaluate an upper bound of edit distance against any overlapping *α*-long path if we choose to drop all variants at *v*. Let Δ_*ins*_, Δ_*del*_ be the longest insertion and deletion variants at vertex *v* respectively. Dropping all variants at *v* can contribute an edit distance of at most Δ_*ins*_ + Δ_*del*_. In cases where only a subset of variant types are present, the bound can be adjusted easily. The following greedy algorithm is designed to select an appropriate set of coordinates where variants can be removed while ensuring that the graph remains (*α, δ*)_*e*_-compatible.

As before, let *p*_1_, *p*_2_, …, *p*_*n*_ be the *n* variant coordinates in *G* in ascending order. Note that an *α*-long path in graph *G* can span *> α* range along the coordinate axis by using deletion edges. For a variant position *p*_*i*_, consider the left-most position *p*_*j*_ such that 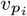can be reached from 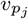by using any path that uses *< α* labeled edges. The rationale for choosing *p*_*j*_ this way is that any *α*-long path which begins at a variant coordinate vertex prior to 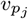cannot pass through vertex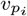. Such a window is pre-computed for each variant position, and we ensure that dropped variants within each window collectively contribute to edit distance ≤ *δ*. To achieve this, our greedy heuristic is to consider the variant positions from left to right. A variant position is removed if and only if the total sum of differences within its window remains ≤ *δ*. It is straightforward to prove that the resulting reduced graph remains (*α, δ*)_*e*_-compatible.

##### Run-time complexity

Computing window lengths for each coordinate vertex is the most time-consuming step in the above algorithm because the remaining steps have linear complexity either in terms of count of variants or count of variant positions in graph *G*. For calculating window lengths in the above algorithm, we can ignore SNP and insertion variants from *G*, and consider only deletion variants. If y denotes the count of deletion variants, then the modified graph will have exactly |R | + 1 coordinate vertices and |R1| + y edges. Any vertex *v*_*i*_ (*i* > 0) has exactly one incoming labeled edge (say, from vertex 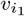) and ≥ 0 incoming unlabeled edges (say, from vertices *v*_*i*2_, *v*_*i*3_,…,*v*_*i*k_). Let the function *f*(*v, x*) indicate the left-most vertex that can be reached from v by using a path that uses < x labeled edges. Then, *f*(*v_i_*, *α*) equals the left-most vertex among 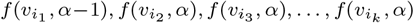. One way to compute this recursion is to compute a vector of values 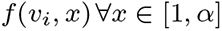for each vertex along the coordinate axis going from left to right. This procedure requires *O* (*α* • (|*R*1| + *y*)) time. In practice, *y* ≪|R1|, so this procedure effectively requires O(α • |R1|) time.

#### 3.2.2 An ILP-based algorithm

Alternatively, we can further improve the above heuristic by using ILP. This can be achieved by formulating the edit distance constraints discussed above for each window as a set of ILP constraints. Similar to our LP-based algorithm for Problem 2, *A* is an *n* × *n* matrix, where row *i* contains non-zero values for those variants that are within the pre-computed window of variant *i*. For instance, if coordinate *p*_*j*_ is within the pre-computed window span of coordinate *p*_*i*_ (*j* ≤ *i*), then *A*[*i*][*j*] is set to the estimated upper bound of differences induced by removing all variants at coordinate *p*_*j*_ as discussed before. Variable *X* is an *n* × 1 boolean column vector, where *X*[*i*] = 1 iff variants at coordinate *p*_*i*_ are removed. Then, ILP constraints required to ensure (*α, δ*)_*e*_-compatibility can be specified as *A* · *X* ≤ *B*, where *B* is a column vector with each value = *δ*. Define *C* to be an *n* × 1 column vector. While addressing Problem 3, set *C*[*i*]’s as 1, and for Problem 4, set 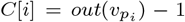 i.e., the count of variants at coordinate *p*_*i*_. In both cases, the ILP objective is set to maximize *C*^*T*^ *X*. These ILP formulations have higher run-time complexity when compared to the greedy solution, but are guaranteed to provide at least as good and possibly superior reduction for both Problems 3 and 4. Neither algorithm guarantees optimality.

## 4 Haplotype-aware variant selection

In the previous problem versions, we considered all *α*-long paths in graph *G*. Here we address the important special case where paths are *restricted* to correspond to strings observed in haplotypes *R*_1_, *R*_2_, …, *R*_*m*_. Due to this restriction, fewer *a*-long strings are checked for mappability. As a result, solutions to the previous problems are sub-optimal for this case because further reduction may be possible. We start by making the simplifying assumption that the input haplotypes contain only SNPs, and have equal length. Outgoing edge label(s) from a coordinate vertex in a reduced graph can come from any of the *m* haplotypes. Graph *G′* is said to be 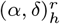 compatible if all *α* -long *restricted* paths in *G* map to *G′* with Hamming distance ≤ *δ* between the corresponding strings. In this scenario, consider the following problems

Problem 5. *Compute an* 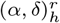*-compatible reduced variation graph G′ with minimum number of coordinates containing one or more variants*.

Problem 6. *Compute an* 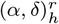*-compatible reduced variation graph G′with minimum number of variants*.

We prove that solving the above problems is 𝒩 𝒫-hard. We give two reductions for Problem 5. The first is a general reduction whereas the second proves hardness for even *δ* = 1. These reductions trivially generalize to Problem 6. Consider the following decision version of Problem 5. Does there exist an 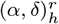-compatible simplified graph *G′* with ≤ *k* coordinates containing one or more variants?

### Lemma 4.

*The decision version of Problem 5 is* 𝒩 𝒫*-complete*.

### Proof.

Clearly, the problem is in 𝒩 𝒫. Recall the decision version of the closest string problem (CSP) (Lanctot *et al*., 2003). Given a set S of strings each of length *l* and a parameter *d*, CSP checks existence of a string that is within Hamming distance of *d* to each of the given strings. CSP is known to be 𝒩 𝒫-complete. CSP exhibits a trivial reduction to Problem 5: Assume the collection of reference haplotypes to be S. The following statements are equivalent: (i) there exists a string with Hamming distance ≤ *d* to each of the given strings in S, (ii) there exists an 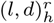-compatible graph *G′* with no coordinate containing one or more variants. As a result, Problem 5, which is stated for an arbitrary value of *k*, is 𝒩 𝒫-complete.

CSP is known to be 𝒩 𝒫-complete even for a binary alphabet, thus also making Problem 5 𝒩 𝒫-complete for a binary alphabet. However, CSP is fixed-parameter tractable relative to parameter *d* (Gramm *et al*., 2003). Consequently, the above claim does not resolve the complexity of Problem 5 for a constant *δ*. For practical applications, *δ* is expected to be small. We address this in the following lemma.

### Lemma 5.

*The decision version of Problem 5 is* 𝒩 𝒫*-complete even if δ* = 1.

### Proof.

Recall the decision version of the maximum independent set (MIS) problem. Given an undirected graph, the MIS problem asks for a set of ≥ *k* vertices no two of which are adjacent. In an MIS graph instance *G*_*m*_ (*V*_*m*_, *E*_*m*_), let 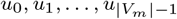 be the vertices and 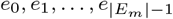 be the edges. We translate this into a multigraph instance of Problem 5 as follows. Let Σ = {A, C}. Define haplotype reference sequences 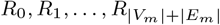 each of length |*V*_*m*_|. The first |*E*_*m*_| sequences are defined using the MIS graph instance *G*_*m*_ while the rest are auxiliary:

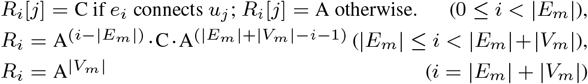

Observe that edge-labeled variation graph *G* built by using the above sequences has a coordinate axis with |*V*_*m*_| + 1 vertices (Figure 3). Each coordinate ∈ [0, |*V*_*m*_|) has two SNPs ‘A’ and ‘C’. *Claim: There exists a k-sized independent set in G*_*m*_(*V*_*m*_, *ϵ*_*m*_) *if and only if there exists* 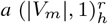*-compatible reduced variation with* |*V*_*m*_| − *k coordinates containing one or more variants*. Consider the forward direction. Suppose there are *k* vertices in an independent set ℐ. Build a reduced variation graph *G′* by removing ‘C’-labeled outgoing edges from coordinates *j* (∀*j*) such that *u*_*j*_ ∈ ℐ. Note that *G′* is 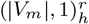 -compatible. Next consider the backward direction. Suppose *p*_1_, *p*_2_, …, *p*_*k*_ are the *k* coordinates in a compatible simplified graph *G′* where variants are removed. If *k* = 1, then finding an independent set ℐof size 1 is trivial. If *k >* 1, then we note that each of the *k* coordinates must have a single outgoing edge labeled with ‘A’ to ensure 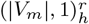-compatibility with respect to the auxiliary reference sequences. It can be further deduced that 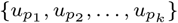 is an independent set of graph *G*_*m*_.

**Fig 3.**
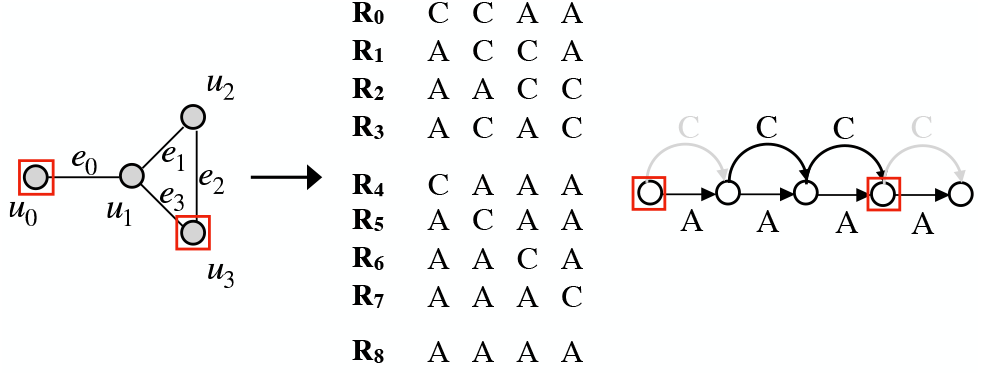
Illustration of reduction used to prove Lemma 5. Vertices selected as independent set are highlighted in red (left). Accordingly, we can find an equivalent reduced variation graph where variants from two vertices are removed (removed edges are highlighted in gray).

## 5 Experimental results

### Hardware and software

We provide C++ implementations of all the algorithms presented in Section 3 (https://github.com/ac-gt/VF). Among these, the first two handle SNP-based variation graphs (*Greedy*_*s*_ and *LP*_*s*_), and the remaining (*Greedy*_*i*_,ILP_*i*_^*v*^ and ILP_*i*_^*v*^ are designed for a generic variation graph containing both SNPs and indels. Our ILP algorithm (Section 3.2.2) supports two different objective functions, the first minimizes count of variants, and the second minimizes variant-containing positions. Accordingly, their naming, i.e., *ILP*_*i*_^*v*^ and *ILP*_*i*_^*p*^ differentiates the two versions respectively.

Using human variation graphs (Table 1), we assess the graph size reduction achieved by the various algorithms, and also evaluate their run-time performance and scalability. The *LP*_*s*_, ILP_*i*_^*v*^ and ILP_*i*_^*p*^ algorithms make use of Gurobi 9.1.0 solver for LP optimization. All the algorithms were tested on dual Intel Xeon Gold 6226 CPUs (2.70 GHz) processors equipped with 2×12 physical cores and 384 GB RAM. Among the implemented algorithms, only the *LP*_*s*_, ILP_*i*_^*v*^ and ILP_*i*_^*p*^ take advantage of multiple cores via Gurobi, whereas the remaining two are sequential.

**Table 1.** Variation graphs used for testing the proposed variant selection algorithms.

| Graph label | Chr | Type of variants | No. of variants | No. of variant containing loci |
| --- | --- | --- | --- | --- |
| vg_chr1_SNP | 1 | SNPs | 6,234,054 | 6,215,039 |
| vg_chr22_SNP | 22 | SNPs | 1,063,618 | 1,059,517 |
| vg_chr1_SV | 1 | Indel SVs | 6,525 | 6,369 |
| vg_chr22_SV | 22 | Indel SVs | 2,056 | 1,996 |

### Variation graph constructio

We tested our algorithms using variation graphs associated with human chromosome 1 (249 Mbp) and chromosome 22 (51 Mbp) respectively. For both chromosomes, we built two types of variation graphs, corresponding to SNPs and indel structural variants (SVs), respectively. Rather than using small-sized indels, we intentionally chose to experiment using indel SVs (≥ 50bp). This is useful to contrast output quality while exploring variant types from point mutations (SNPs) to larger variants. Exclusion of SVs is naturally expected to introduce more differences in *α*-sized paths, and therefore SVs test the limits of our algorithms. SNP variants were downloaded from the 1000 Genomes Project Phase 3 (Consortium *et al*., 2015), and indel SVs were downloaded from a recent long-read based SV survey of fifteen diverse human genomes (Audano *et al*., 2019). We used vcftools (Danecek *et al*., 2011) to pre-process and extract SNPs from the 1000 Genomes Project variant files. Similarly, SVs other than insertions or deletions were filtered out from the SV files. Summary statistics of these variants, and graphs built using them, are listed in Table 1.

### *α* and *δ* parameters

We tested our algorithms using *α* values of 150 bp, 1 kbp, 5 kbp, and 10 kbp. The first is useful for Illumina reads, whereas the latter are useful for different protocols available for long read sequencing (e.g., DNA or RNA sequencing using either PacBio or ONT). For each *α* value, we experimented with *δ* values that are 1%, 5% and 10% of *α*. Here 1% corresponds to low error-tolerance of a mapping algorithm, and 10% corresponds to significant tolerance.

### Performance of *Greedy*_*s*_ and *LP*_*s*_ algorithms

These algorithms were tested using vg_chr1_SNP and vg_chr22_SNP (Figure 4) graphs. For both the algorithms, we report four statistics: (i) count of variants retained, (ii) count of variant-containing loci retained, (iii) run-time, and (iv) peak memory-usage. *LP*_*s*_ and *Greedy*_*s*_ algorithms are guaranteed to return optimal graphs in terms of the objectives (i) and (ii) respectively. The results in Figure 4 suggest that the two algorithms perform almost equally well in terms of both objectives for all tested combinations of (*α, δ*) values. Increasing *α* value while keeping *δ* as a constant fraction of *α* naturally corresponds to fewer SNPs retained. The same is true when *δ* is increased while keeping *α* fixed. These results corroborate the fact that longer reads and higher sensitivity of mapping algorithms result in retention of fewer variants in a variation graph. For instance with (*α* = 10 kbp, *δ* = 1000), the *Greedy*_*s*_ algorithm retained only 531 (0.009%) out of 6,234,046 SNPs using graph vg_chr1_SNP. After this run, average distance between two adjacent loci containing SNPs increased from 39 to 225, 963. This suggests that variation selection algorithms can potentially yield long stretches of variant-free regions in graph, where the usual read-to-sequence mapping algorithms can also operate.

**Fig 4.**
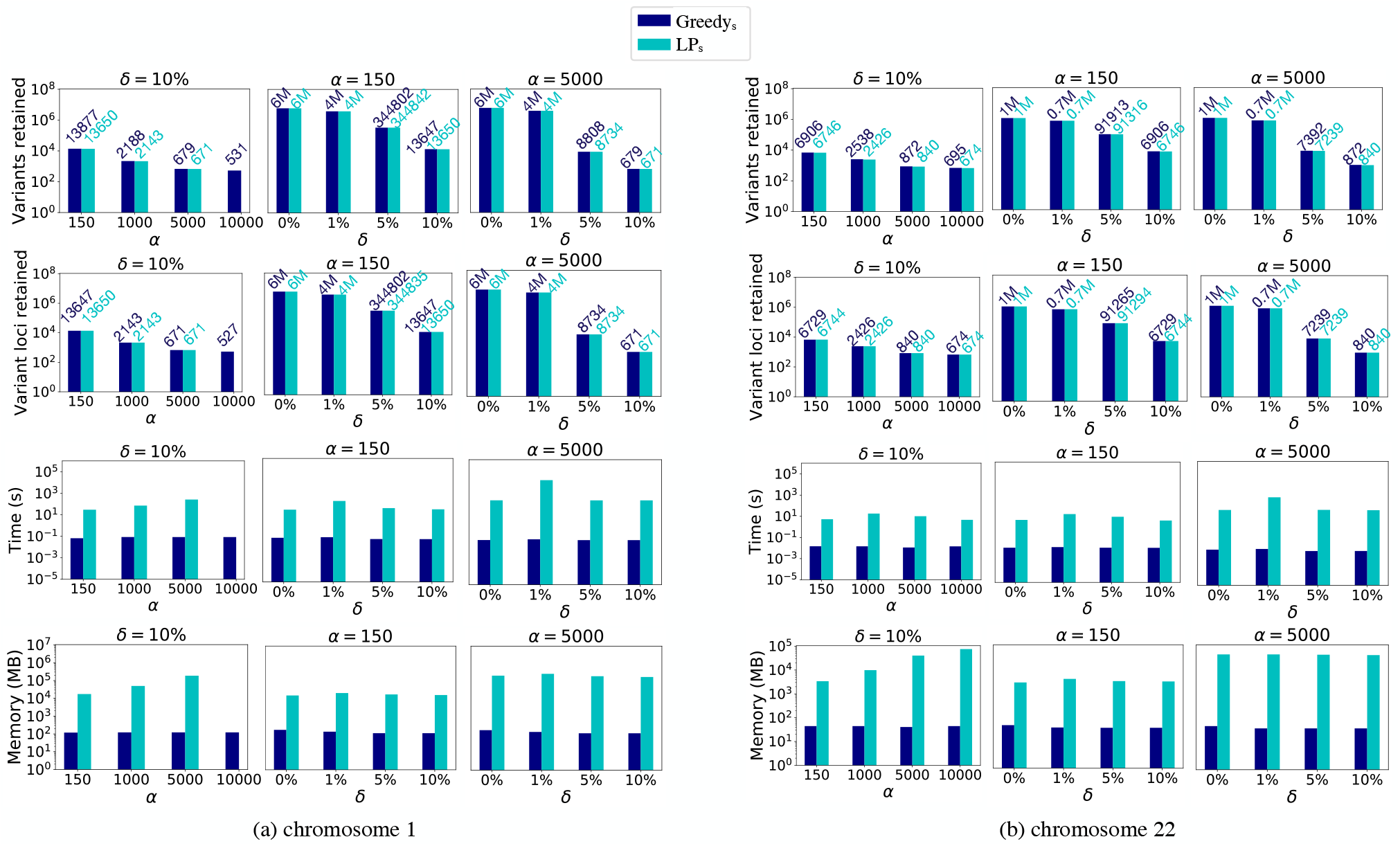
Empirical evaluation of Greedys and LPs algorithms using two human variation graphs vg_chr1_SNP and vg_chr22_SNP containing SNPs. These plots demonstrate reduction achieved in graph sizes while varying *α* and δ parameters. Size of the complete variation graph (*δ* = 0%) is included for comparison. Numbers on top of bars present actual data, useful for comparison when both Greedys and LPs achieve close results. Result of LPs algorithm is missing for *α* = 10, 000 (left-most column) because Gurobi LP solver crashed due to insufficient memory. Y-axes are log-scaled in all the above plots.

If run-time is considered, the *Greedy*_*s*_ algorithm runs significantly faster than *LP*_*s*_, which was also reflected by our time complexity analysis in Section 3.1. Using graph vg_chr1_SNP, *LP*_*s*_ algorithm ran out of memory for *α* = 10 kbp due to increased size of the matrix, i.e., count of non-zeros in the matrix which specifies the LP constraints. Taken together, *Greedy*_*s*_ algorithm suffices for most practical purposes because it is fast and optimal in terms of its objective to minimize count of variant-containing loci retained. *Greedy*_*s*_ algorithm does not account for the number of variants at a locus while deciding its fate, yet it achieves near-optimal reduction in terms of minimizing the count of variants. This is likely because most human SNPs are biallelic.

### Performance of *Greedy*_*i*_ and ILP-based algorithms

We tested our *Greedy*_*i*_, ILP_*i*_^*v*^ and ILP_*i*_^*p*^ heuristics using vg_chr1_SV and vg_chr22_SV (Figure 5) graphs. In contrast to SNPs which are single-base mutations, the sizes of SV indels in chromosome 1 computed by Audano *et al*. (2019) range from 50 bp to 33 kbp, with mean length 0.5 kbp. As a result, it is natural to expect that the fraction of variants retained will be much higher compared to SNPs. The ILP-based heuristics are guaranteed to achieve superior results than the greedy heuristic, i.e., ILP_*i*_^*v*^ heuristic is expected to retain the smallest count of variants among the three, and similarly *ILP*_*i*_^*p*^ heuristic will retain the smallest count of variant-containing loci. For instance, with long-read compatible settings (*α* = 10 kbp, *δ* = 1000), the ILP_*i*_^*v*^ and ILP_*i*_^*v*^ *Greedy*_*i*_ heuristics retained 26.8%, 27.0% and 31.3% SVs respectively in graph vg_chr1_SV. Similarly, 24.8%, 25.0% and 30.6% SVs were retained in graph vg_chr22_SV. However, with short-read compatible settings (*α* = 150 bp, *δ* = 8), all SVs were retained by all three heuristics, as expected. These results are not necessarily optimal, but we do not expect them to deviate significantly from optimal numbers. The rationale is that not only SVs are bigger in size but also SV loci are known to be clustered in several known hot-spots of the human genome, e.g., within the last 5 Mbp of both chromosome arms (Audano *et al*., 2019).

In terms of count, SVs occur much less frequently as compared to SNPs or indels. As a result, run-time of all the three heuristics was dominated by their first step of computing the constraints required to ensure (*α, δ*)_*e*_ compatibility, which is common in all of them. As shown in Section 3.2, this step requires time proportional to *α* as well as the length of the reference sequence. Accordingly, we observe that running time is comparable among all three heuristics, appears to be independent of *δ*, scales roughly linearly with *α*, and time spent is higher using graph vg_chr1_SV than vg_chr22_SV. With the largest *α* = 10 kbp value, all three algorithms require about six minutes and one minute to process the two graphs, respectively.

**Fig 5.**
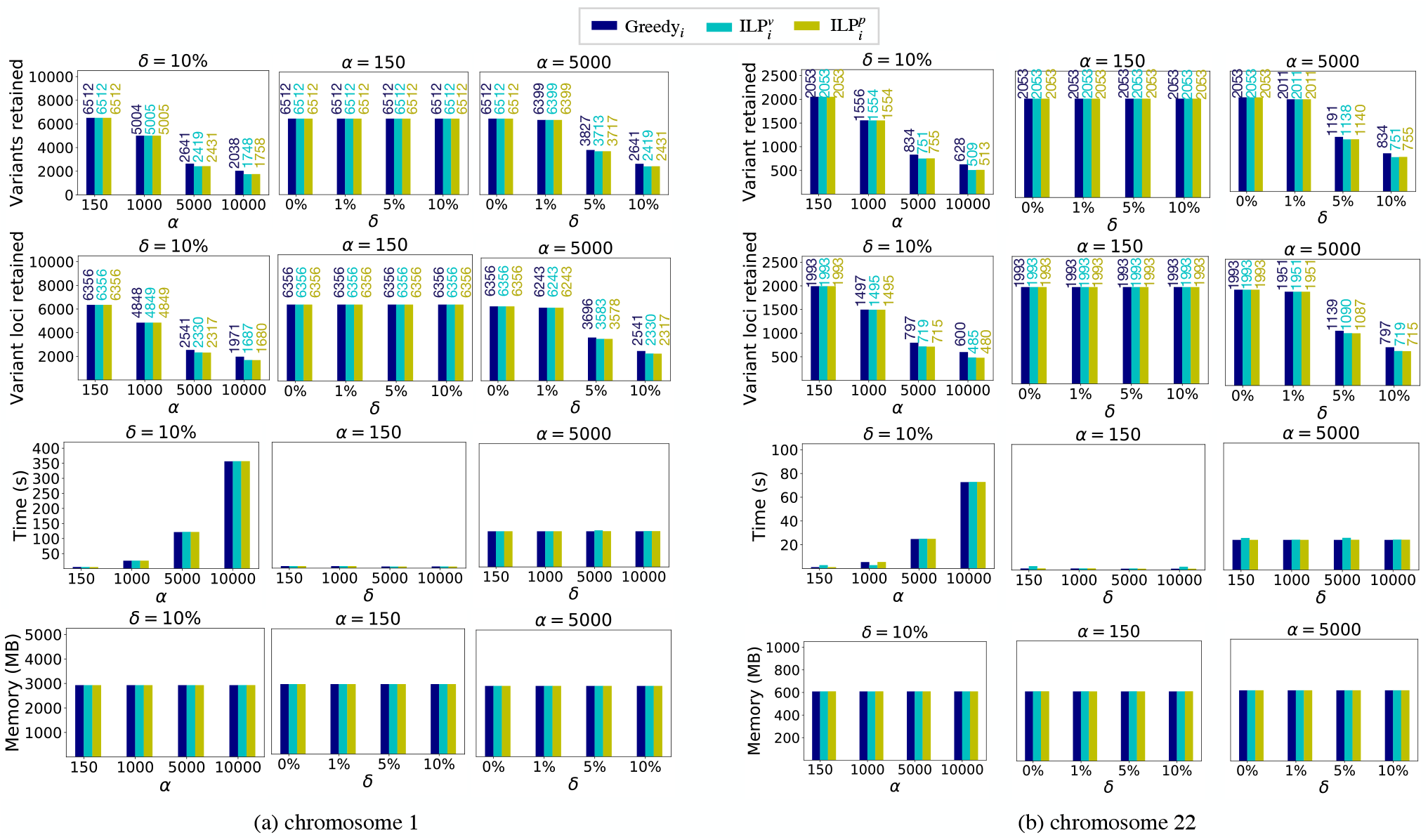
Empirical evaluation of Greedyi, ILP_*i*_^*v*^ and ILP_*i*_^*v*^ algorithms using two human variation graphs vg_chr1_SV and vg_chr1_SV containing insertion and deletion structural variants. These plots demonstrate reduction achieved in graph sizes while varying *α* and *δ* parameters. Numbers on top of bars present actual data, useful for comparison when the three algorithms achieve close results.

### Impact on sequence-to-graph mappers

We conducted a preliminary evaluation to assess the impact of variant selection on read-to-graph mapping. For this experiment, we considered the reduced graph obtained by ILP_*i*_^*v*^ using vg_chr1_SV graph as input. As discussed previously, the *ILP*_*iv*_ heuristic retained 1, 748 of the 6, 512 SVs using *α* = 10 kbp and *δ* = 1000 parameters. We built two variation graphs using VG (v1.29.0) tool corresponding to the complete set of SVs and the reduced set of SVs respectively. The graph statistics (e.g., vertex degree distribution) were validated to ensure the presence of SVs in the respective graphs. Next, we simulated 10, 000 long reads, each of length 10 kbp from randomly chosen paths of the complete variation graph with error-rate 5% using VG’s read simulation feature. Subsequently, we made use of GraphAligner (v1.0.11) to map these reads to both variation graphs.

We observed the following. First, all 10, 000 reads were successfully mapped by GraphAligner to both the graphs. Second, each read was mapped only using primary alignments, and there were no secondary alignments reported. This indicates that there was no mapping ambiguity while using the reduced variation graph. Finally, the impact of missing variations was observed in the count of reads with split-read alignments, which increased from 2 to 209. Split read alignments are often used as a signature by variant callers to discover SVs (Rausch *et al*., 2012). A direct comparison of mapping coordinates could not be done because VG used different vertex identifiers in the two graphs which have different topology. We note that GraphAligner required similar runtime and memory in both scenarios (about five minutes). This is likely because the graph is nearly linear, due to limited count of SVs (6, 512) that were available in the complete chromosome 1 graph. This result is preliminary, but motivates a deeper investigation into the impact of the proposed algorithms on various sequence-to-graph mapping algorithms while using a much larger catalog of variants as input. Variant selection tool FORGe (Pritt *et al*., 2018) uses allelic frequency data as an input to its model. Currently, frequency assessment remains challenging in case of SVs due to the lack of appropriate tools as well as data (Mahmoud *et al*., 2019). A direct comparison with FORGe could not be carried out due to these limitations.

## 6 Conclusions and open problems

We developed a novel mathematical framework for variant selection, and presented multiple algorithms and complexity results for various problems arising from this framework. Experimental results demonstrate substantial reduction in the resulting variation graph sizes, while guaranteeing bounds on the number of errors tolerated while doing so. Implementations of all the four algorithms that are proposed in this paper are available as open-source, and can be used by practitioners for downstream pan-genomic analysis. The path-length and error parameters (*α, δ*) can be tuned to match the choice of sequencing technology, mapping algorithms, and types of variants considered. We demonstrated that a high fraction of small-scale variants, but no large-scale variants can be left out during short-read mapping to variation graphs. On the other hand, almost all small-scale variants and a significant fraction of large-scale variants may be excluded prior to long-read-based analysis.

The proposed variant selection framework underpins a rich class of problems making it fertile ground for future research. (i) Optimal algorithms for the two problems associated with indel variants (Problems 3 and 4) are unknown. In fact, it is not known whether these two problems can be solved in polynomial time. (ii) While we were able to prove that the haplotype-aware versions of the problem are 𝒩 𝒫-hard, efficient heuristics and approximation algorithms for these problem are yet to be developed. Haplotype-aware algorithmic extensions can result in further reduction of graph sizes because fewer paths need to be preserved. (iii) It may also be possible to further extend this framework, and add constraints similar to allele frequency thresholding, e.g., by asking a reduced graph which is allowed to violate error-bound for up to 1% of haplotypes at any position. (iv) Finally, this work only considered pre-dominant variant categories -SNPs, insertions and deletions; further research is needed to analyze other variant types such as duplications, inversions, and complex genomic rearrangements.

